# Heterotypic interaction promotes asymmetric division of human hematopoietic stem and progenitor cells

**DOI:** 10.1101/2023.08.31.555706

**Authors:** Adrian Candelas, Benoit Vianay, Matthieu Gelin, Lionel Faivre, Jerome Larghero, Laurent Blanchoin, Manuel Théry, Stephane Brunet

## Abstract

Hematopoietic Stem and Progenitor Cells (HSPCs) give rise to all cell types of the hematopoietic system through various processes including asymmetric divisions. However, the contribution of stromal cells of the hematopoietic niches in the control of HSPC asymmetric divisions remains unknown. Using polyacrylamide microwells as minimalist niches, we show that heterotypic interaction with osteoblast promotes asymmetric division of human HSPC. Upon interaction, HSPCs polarize in interphase with centrosome, the Golgi apparatus and lysosomes positioned close to the site of contact. Subsequently during mitosis, HSPCs orient their spindle perpendicular to the plane of contact. This division gives rise to siblings with unequal amounts of lysosomes and differentiation markers such as CD34. Such asymmetric inheritance generates heterogeneity in the progeny, which is likely to be a key contributor to the plasticity of the early steps of hematopoiesis.

## Introduction

Asymmetric cell division is one evolutionary conserved strategy to generate cellular diversity adopted by stem cells. Asymmetric divisions can lead to unequal inheritance of cell fate determinants ^1^ or place the two daughter cells in distinct environments providing different fate determinants ^2^. In most characterized stem cells so far, asymmetry relies on the polarization of the mother cell controlled by external cues of the cell microenvironment ^3 4 5^. These cues induce centrosome positioning in interphase, and in turn accurate orientation of the mitotic spindle along the polarization axis ^6^. Paradigmatic of adult stem cells, Hematopoietic Stem and Progenitor Cells (HSPCs) self-renew and differentiate into lineage-determined daughter cells, that will give rise to all cell types of the hematopoietic system ^7^. HSPCs are retained in the bone marrow within specific niches ^8^, where external cues modulate their homeostasis ^9 10^. However, the mechanism leading to asymmetric division of HSPCs has not yet been well characterized. *In vitro* HSPCs can undergo asymmetric divisions ^11 12 13 14 15 16 17^. In addition, investigations conducted *in vitro* ^18 19 20 21^ or *in vivo* ^22 23 24 25^ have shown that stromal cells including osteoblasts can modulate HSPCs proliferation and differentiation. This suggests that stromal cells may play a role in controlling HSPC asymmetric divisions ^26^. Nevertheless, such a role still remains hypothetical. Classical co-culture systems are unlikely to allow deciphering the mechanisms at play since they scantily recapitulate the long-term confinement encountered by HSPCs in their niche. Moreover, *in vivo,* imaging of HSPC-stromal cell interactions and the impact on HSPC divisions has not yet been achieved. To overcome such limitations, systems of intermediate complexity have been developed ^27 28 29 30 31^. We have recently set-up a system of microwells as “minimalist niches” to show that human HSPCs in interphase have the capacity to interact with specific stromal cells of the bone marrow and polarize by reorganizing intensively their intracellular architecture ^32^. Using this system, we here investigate the impact of heterotypic interactions on the mode of division of HSPCs.

## Results

### Osteoblast interaction induces stable HSPC polarization in interphase

To overcome limitations of classical co-culture for long-term tacking of interactive HSPCs (Figure S1A), polyacrylamide microwells were used. They were either coated with fibronectin, a major extracellular matrix protein in the bone marrow ^33 34^ or seeded with osteoblast as a stromal cell of the bone marrow, or skin fibroblast as a control cell absent from the hematopoietic niches (Figure 1A, Figure S1). HSPCs were first labelled with fluorescent CellTracker at concentration optimized to have no impact on cell viability, cell cycle progression and proliferation (Figure S1D-G) and then loaded at a density of one cell per well (Figure 1A and B, Figure S1B and C). Cell-substrate adhesion was restricted to the bottom of the microwell, preventing cell escape and migration (Figure S1A). Using this system, cell division could be tracked over few generations (Figure 1B).

**Figure 1.**
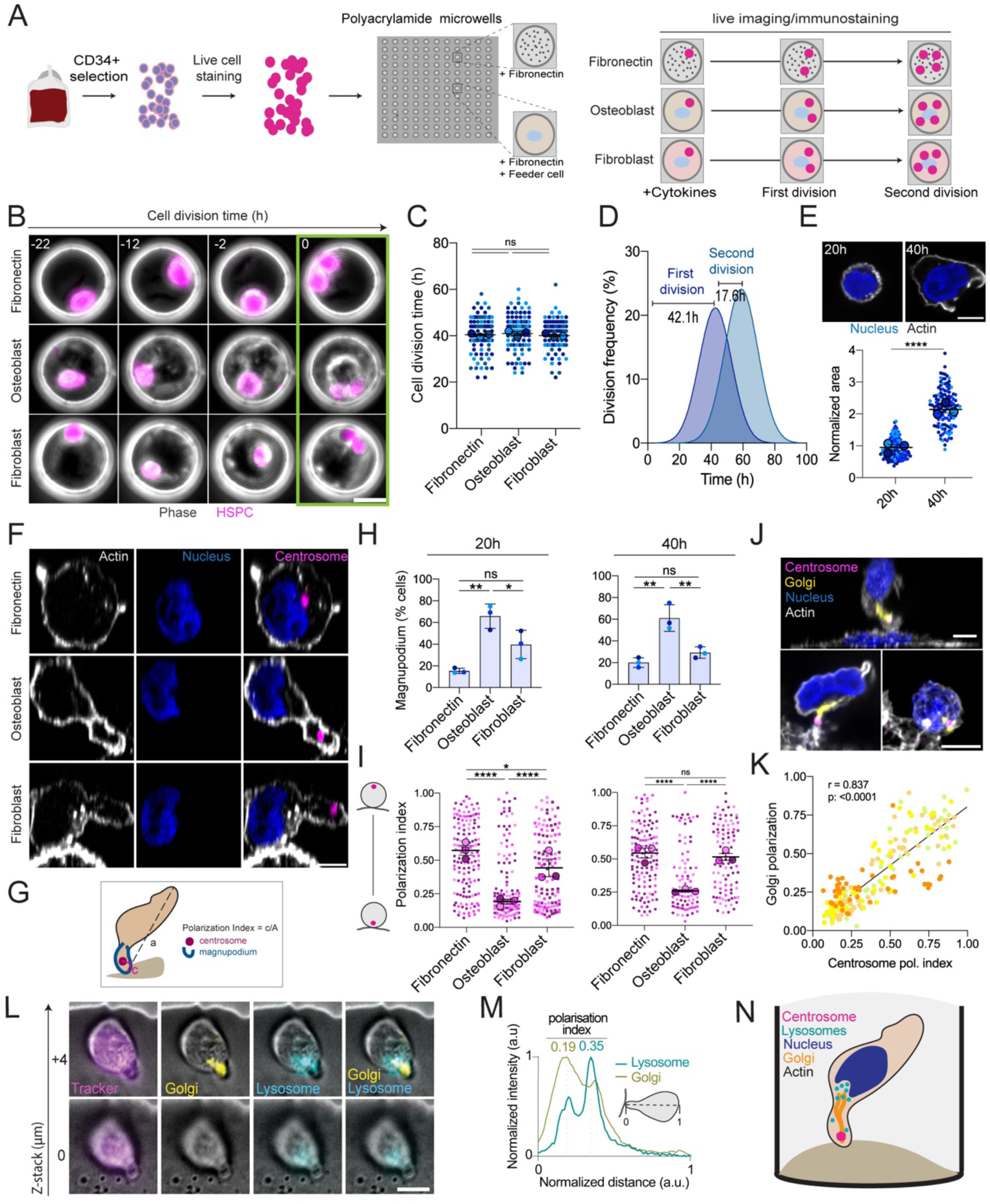
Osteoblast interaction induces stable HSPC polarization in interphase. **(A)** Schematic representation of the experimental design for long-term tracking of HSPCs (co-)cultured in polyacrylamide microwells. **(B)** Time-lapse monitoring with transmitted light and fluorescence of live HSPCs labelled with CellTracker (magenta) in contact with fibronectin (upper row), osteoblasts (middle row), and skin fibroblasts (lower row). Time is indicated in hours relative to HSPC mitosis (framed in green). Scale bar: 20µm. **(C)** Superplot of the cell division time of HSPCs in contact with fibronectin, osteoblast, and skin fibroblast. Each color represents a biological replicate (three biological replicates; fibronectin n_tot_=97, osteoblasts n_tot_=95, fibroblasts n_tot_=77 cells). For each replicate, the median value appears as a large disk of the corresponding color. Mean of medians ± SEM shown as black bars. One-way ANOVA test. n.s: non-significant. **(D)** Gaussian fitted curves of the relative frequency for the first and second cell division (n_tot_=122, (n_tot_=141 cells respectively; three biological replicates). The peaks of division are indicated. **(E)** Representative images of HSPCs on fibronectin. Actin is shown in gray, nucleus in blue. Scale bar: 5µm. On the bottom, SuperPlot of the HSPC normalized area analyzed upon culture on fibronectin at 20h and 40h of culture. Each color represents a biological replicate (three biological replicates; 20h n _cells_= 141, 40h n _cells_= 121). For each replicate, the median value appears as a large disk of the corresponding color. Mean of medians ± SEM shown as black bars. Mann-Whitney U test. ****: P≤0.0001. **(F)** Representative images of HSPC in contact with fibronectin, osteoblasts, and skin fibroblasts after 40h of culture. Actin is shown in gray, nucleus in blue and the centrosome in magenta. Scale bar: 5 µm. **(G)** As schematized, the HSPC polarization index was defined as the ratio between *c,* the distance from the site of contact to the projected centrosome point, and *a,* the HSPC length. **(H)** Percentage of HSPC with a magnupodium 20h (left) and 40h (right) of culture (three biological replicates). One-way ANOVA test. n.s : non-significant; *: P ≤0.05; **: P≤0.01; **(I)** SuperPlot of HSPC polarization index at 20h of culture (left panel; three biological replicates: fibronectin n_tot_=142 cells, osteoblasts n_tot_ =142 cells, skin fibroblasts n_tot_ =125 cells) and at 40h of culture (three biological replicates: fibronectin n_tot_=127, osteoblasts n_tot_=91, skin fibroblasts n_tot_ =96 cells). For each replicate, the median value appears as a large disk of the corresponding color. Mean of medians ± SEM shown as black bars. Kruskal–Wallis ANOVA test. ns: non-significant; *: P ≤0.05; ****: P≤0.0001. **(J)** Representative images of fixed HSPC polarizing (upper and lower left images) or unpolarized (lower right image) in contact with osteoblasts after 20h of culture. Centrosome appears in magenta, Golgi in yellow, nucleus in blue and actin in white. Scale bar = 5µm. **(K)** Correlation analysis between polarization index and Golgi polarization after 2h (in green), 4h (in light orange), 8h (in yellow), and 16h (in dark orange) of culture on osteoblast (one biological replicate, n_tot_=219 cells). Spearman correlation test is indicated. **(L)** Representative transmitted light and fluorescence time-lapse images, at two 4µm distant Z-stacks, of live HSPCs in contact with osteoblast. HSPC is labelled with CellTracker (magenta), Golgi tracker (yellow), and LysoBrite (Cyan) for lysosome live staining. Scale bar: 5 µm. **(M)** Representative line-scan analysis of a polarized HPSC for lysosome and Golgi signal intensities along the cell axis (as schematized in grey). **(N)** Schematic representation of HSPC polarization upon interaction with osteoblast.

Cells enlarged between 20 and 40 h of culture indicative of cell cycle progression (Figure 1E) and the first cell division took place around 40h after cytokine addition with no significant difference in the three culture conditions (Figure 1C). The first HSPC cell cycle was extended compared to the next generation cell cycle (Figure 1D). This result indicates that feeder cells present in the microwells had no effect on HSPC cell cycle duration and division kinetics.

HSPCs cultured on fibronectin or in presence of skin fibroblast mostly exhibited a round morphology or presented a uropod on their back after 20h and 40 h of culture (Figure 1F and Figure S2A). In contrast, around 60% of HSPCs contacting osteoblasts were strongly elongated exhibiting a magnupodium anchored at the site of contact, as previously described ^32^. The magnupodium, present at 20h of culture, was still observed after 40h prior to cell division (Figure 1F-H, Figure S2A).

Magnupodium formation is associated with centrosome positioning at its tip, in close proximity of the contact site with the stromal cell ^35 32^. In HSPCs interacting with osteoblast, the centrosome was indeed located at the tip of the magnupodium at 20h and 40h of culture (Figure 1F and I, Figure S2A). Moreover, the cell polarity index, defined as the ratio between the centrosome distance to the contact site with respect to cell length was significantly lower in HSPCs interacting with osteoblasts compared with HSPCs cultured on fibronectin or seeded on fibroblast at 20h and 40h of culture indicative of a stable polarization (Figure 1I).

Organelles distribution in polarized HSPCs was then examined. Golgi was found to be apposed to the centrosome (Figure 1J and K), elongating within the magnupodium. In addition, using cell permeable and stable fluorogenic markers, lysosomes were found tightly associated with the Golgi in the magnupodium and extending toward the cell body (Figure 1L and M, Figure S2B and C).

These results indicate osteoblast-HSPC interaction induces HSPC polarization marked by the formation of a magnupodium with the centrosome located at the tip toward the site of interaction and Golgi and most of the lysosomes confined within the magnupodium out of the cell body (Figure 1N). Such polarized structure is stably established and maintained during cell cycle progression up to mitosis.

### Osteoblast interaction promotes HSPC perpendicular spindle orientation at mitosis

We then investigated the impact of HSPC polarization on the subsequent mitosis. Since the first division was found to occur around 40h, cells were fixed at this time to enrich mitotic HSPCs population.

At mitosis onset, HSPCs in the three conditions were observed to round-up (Figure 2A). At least one of the two separating centrosomes was found close to the site of contact in 47% of HSPC interacting with osteoblasts, in contrast to HSPC cultured on fibronectin or on fibroblasts (Figure 2A and B). At metaphase and anaphase, spindles were preferentially oriented parallel to the well bottom in HSPC cultured on fibronectin, as previously documented ^15^ (Figure 2E, left panel). Similarly, the spindle was mainly oriented parallel to the surface of contact with fibroblasts (Figure 2E, right panel). In contrast, in 68% of HSPCs interacting with osteoblasts, the metaphase spindle was perpendicular to the cell surface and this orientation was maintained in anaphase (Figure 2E, central panel). These results indicate that the interaction of HSPC with osteoblast specifically promotes spindle positioning perpendicularly to the osteoblast surface.

**Figure 2.**
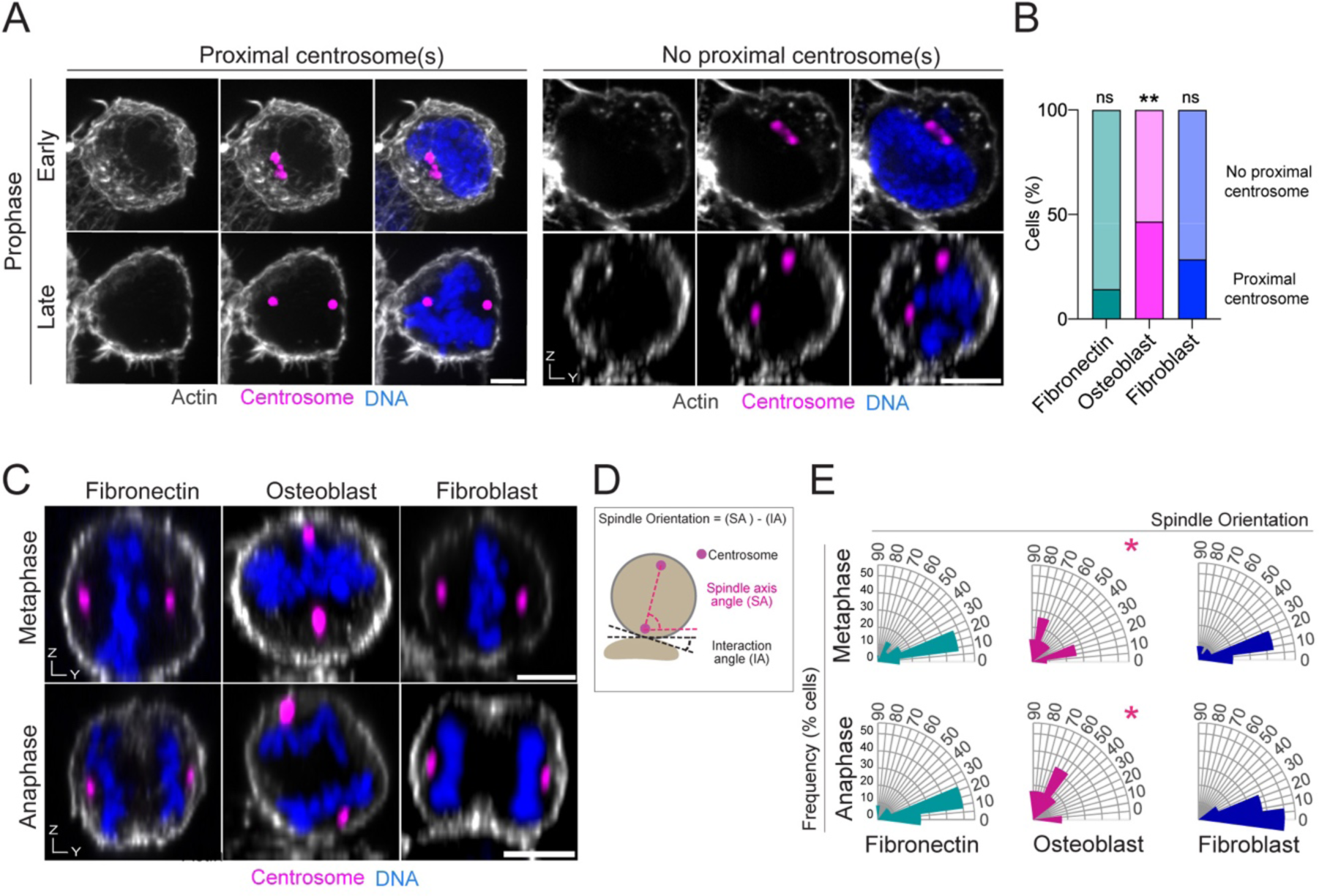
HSPCs divide perpendicularly to the surface of contact with osteoblast. **(A)** Representative images of HSPC in contact with osteoblast at early and late prophase (upper and lower row respectively). Actin appears in gray, DNA in blue, and the centrosome in magenta. Scale bar: 5µm. **(B)** Percentage of HSPCs at prophase exhibiting either proximal centrosome(s) or non-proximal centrosomes (respectively appearing in bright and light colors) upon culture on fibronectin (n_tot_ =21 cells), osteoblast (n_tot_ =15 cells) or skin fibroblast (n_tot_ =8 cells). Binomial test, taking fibronectin as reference result: ns: non-significant; **: P≤0.01 **(C)** Representative images of HSPC at metaphase (upper row) and anaphase (lower row) upon culture in contact with fibronectin, osteoblast, and skin fibroblast. Actin appears in white, DNA in blue and the centrosome in magenta. Scale bar: 5µm. **(D)** As schematized, the spindle orientation angle was calculated as the difference between the spindle axis angle to the horizontal plane and the feeder cell angle (or well bottom) to the horizontal plane. **(E)** Rose diagram representing the frequency of spindle orientation angles in degrees. Upper row: metaphase cells; lower row: anaphase cells. (Metaphase: seven biological replicates; fibronectin n_tot_ =55, osteoblast n_tot_ = 22 and fibroblast n_tot_ = 25 cells; Anaphase: three biological replicates; fibronectin n_tot_ =12, osteoblast n_tot_ = 7 and fibroblast n_tot_ = 9 cells). Kruskal-Wallis ANOVA test. non-significant differences between fibronectin and skin fibroblast for both metaphase and anaphase (P>0.05). In metaphase, *: P ≤ 0.05 for osteoblast vs fibronectin and for osteoblast vs skin fibroblast. In anaphase *: P =0.0266 osteoblast vs skin fibroblast; P =0.0689 osteoblast vs fibronectin.

### Osteoblast-HSPC interaction increases asymmetric inheritance of lysosomes in HSPC daughter cells

In many stem cells or developmental systems, centrosome positioning in interphase according to external cues and subsequent mitotic spindle orientation are hallmarks of asymmetric cell division ^6^. HSPC asymmetric cell division *in vitro* is marked by unequal segregation of lysosomes into the daughter cells ^36^. Therefore, we tested whether heterotypic osteoblast interaction was inducing lysosome asymmetric inheritance. HSPC were labelled with cell permeable and stable fluorogenic CellTracker and LysoBrite^TM^, respectively as cell and lysosome markers. While CellTracker appeared homogeneously segregated in the daughter cells, both symmetric and asymmetric segregation of lysosomes were observed (Figure 3A). The LysoBrite signals were stable in time for both daughter cells (Fig3.B) indicating that the inherited pools were maintained in the daughter cells. The lysosomes inheritance ratio was significantly higher in HSPCs interacting with osteoblasts than the ones cultured on fibronectin or skin fibroblasts (Figure 3C). Setting 1.5 as a threshold value to discriminate symmetric (<1.5) versus asymmetric (≥1.5) inheritance ^17^, we found that asymmetric inheritance was 20-fold higher in the HSPCs interacting with osteoblasts (Figure 3D).

**Figure 3.**
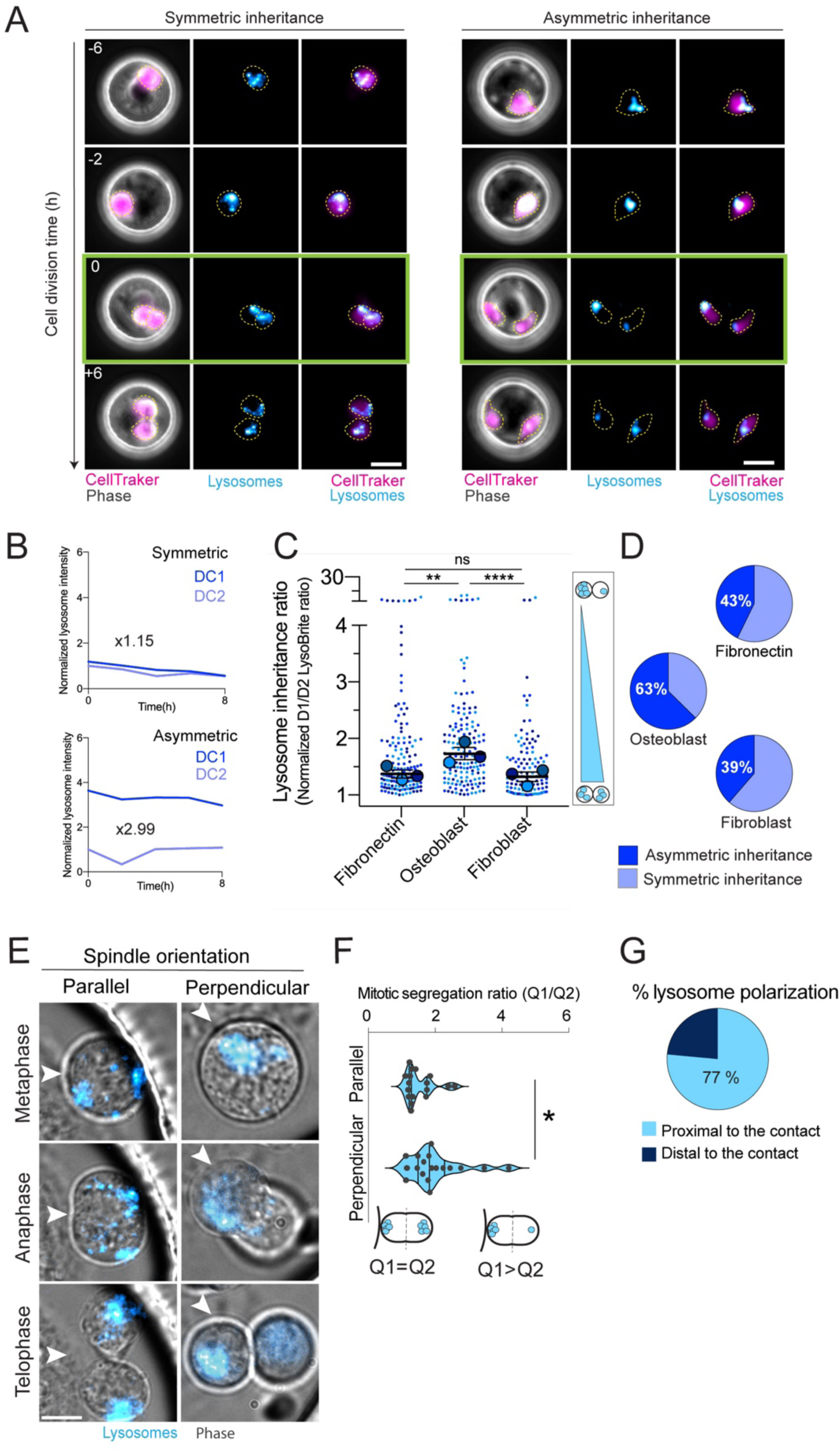
Osteoblast interaction promotes asymmetric lysosome inheritance associated with perpendicular division of HSPC. **(A)** Time-lapse monitoring with transmitted light and fluorescence of live HSPCs on osteoblast, labelled with CellTracker (magenta) and LysoBrite (Cyan). Representative images of LysoBrite symmetric and asymmetric inheritance are presented (left and right panel respectively). Time is indicated in hours relative to cell division, framed in green (t=0). Scale bar: 20µm. **(B)** Analysis of normalized LysoBrite signal intensity in time for the two daughter cells (DC1 and 2) for symmetric and asymmetric (upper and lower graphs) lysosome inheritance. Inheritance ratio mean expressed in xN. **(C)** SuperPlot of normalized LysoBrite inheritance in the daughter cells for HSPCs in contact with fibronectin, osteoblast and skin fibroblast. Each color represents a biological replicate (three biological replicates; fibronectin n_tot_ = 152, osteoblasts n_tot_=158, fibroblasts n_tot_=119 cells). For each replicate, the median value appears as a large disk of the corresponding color. Mean of medians ± SEM shown as black bars. Kruskal–Wallis ANOVA test. ns: non-significant; **: P≤0.01; ****: P≤0.0001. **(D)** Pie chart representation of the distribution of LysoBrite symmetric and asymmetric (light and dark blue respectively) inheritance in HSPC (three biological replicates; fibronectin n_tot_=152, osteoblasts n_tot_=158, skin fibroblasts n_tot_=119 cells). **(E)** Representative time frames of HSPC on osteoblast, taken at metaphase, anaphase and telophase (upper, middle and lower panel), with either parallel (left panel) or perpendicular (right panel) spindle axis orientation. LysoBrite appears in cyan. Arrows highlight the point of contact. Scale bar: 5 µm. **(F)** Violin plot representation of lysosome asymmetry ratio in HSPC according to their spindle axes: parallel or perpendicular (respectively n_tot_=17 and n_tot_=18 cells). Mann-Whitney test. *: P ≤0.05. **(G)** Pie chart representation of the distribution of LysoBrite signal at perpendicular metaphase/anaphase in cell domains proximal (dark blue) and distal (light blue) to the site of contact (n_tot_=17 cells).

Lysosomes segregation was further analyzed during mitosis, according to the cell division geometry. In the case of HSPCs dividing in parallel to the osteoblast surface, lysosomes were equally segregated to opposite cell poles, while in HSPCs dividing perpendicularly, one of the cell poles was enriched in lysosomes (Figure 3E and F). Moreover, most of the lysosomes were found proximal to the site of contact (Figure 3G).

These results show that osteoblast interaction promotes a perpendicular spindle orientation associated with an asymmetric inheritance of lysosomes into HSPC siblings, the proximal daughter cell inheriting most of the mother lysosome pool.

### Interaction with osteoblast leads to HSPC asymmetric division

Is there a causal link between the osteoblast-driven HSPC polarization and the asymmetry of the HSPC division? To address this question, lysosomes distribution and cell division geometry were analyzed using video microscopy in HSPCs interacting with osteoblast, covering late interphase, cell division, and daughter cells interphase (Figure 4A). In parallel, lysosomes polarization indexes were measured in interphase, during mitosis (Figure4C) and in late telophase when daughter cells are individualized (Figure 4D). Based on the distribution of lysosomes and Golgi in polarized HSPCs (Figure 1M), a threshold value of 0.35 was set to discriminate lysosomes polarized versus unpolarized distributions (respectively below and above 0.35).

**Figure 4.**
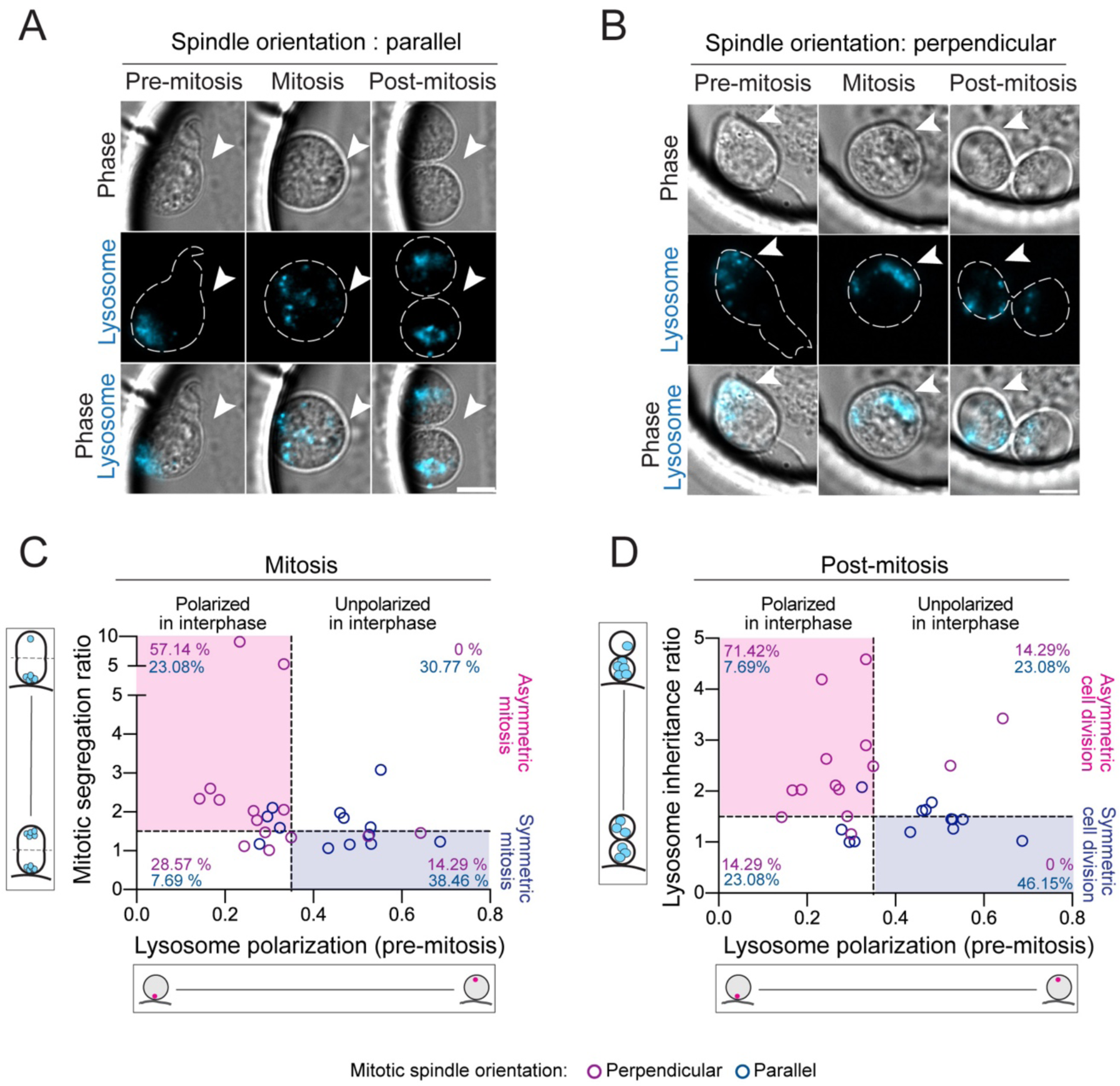
HSPC polarization and perpendicular division precede asymmetric division. **(A)** Time-lapse monitoring with transmitted light and fluorescence (LysoBrite staining appears in Cyan) of live unpolarized HSPCs on osteoblast, dividing parallelly to the surface of contact. Lysosomes are symmetrically segregated. Arrows highlight the point of contact. Cells are contoured using white dashed lines. Scale bar: 5 µm. **(B)** Time-lapse monitoring with transmitted light and fluorescence (LysoBrite staining appears in Cyan) of live HSPCs on osteoblast, dividing perpendicularly to the surface of contact. lysosomes are unequally segregated. Arrows highlight the point of contact. Cells are contoured using white dashed lines. Scale bar: 5 µm. **(C)** Correlation plots representation of the lysosome segregation ratio at metaphase versus lysosome polarization at the interphase of HSPCs in contact with osteoblast (four biological replicates; n_tot_=27 cells). **(D)** Correlation plot representation of lysosome inheritance ratio after cell division, versus lysosome polarization at the interphase preceding mitosis of HSPCs in contact with osteoblast (four biological replicates; n_tot_=27 cells) In C and D, HSPC exhibiting parallel and perpendicular spindle orientation appear respectively in blue and magenta circles, respectively Quadrants limits using 0.35 as threshold a value of lysosome polarization index at interphase to mark polarized and unpolarized HSPCs, and using 1.5 as a threshold value for lysosome asymmetry ratio are presented as dashed lines to emphasize symmetric and asymmetric mitosis.

HSPCs that did not exhibit polarization in interphase were mostly found to divide parallelly to the cell surface (Figure 4A). At mitosis, lysosomes were redistributed within the whole rounded cell (Figure 4A and C, blue box) and were eventually inherited symmetrically in the siblings (Figure 4A and D blue box). In contrast, HSPCs that were polarized in interphase mostly divided perpendicularly to the osteoblast surface (Figure 4B). Lysosomes were clustered in interphase close to the site of interaction in the magnupodium. They remained asymmetrically distributed in mitosis (Figure 4B and C pink box), clustered close to the site of contact (Figure 4B and D, pink box).

These results show that heterotypic interaction with osteoblast is a *bona fide* external cue driving HSPC asymmetric division. Interestingly, not all HSPCs do respond to this cue and some undergo symmetric division. Nevertheless, once the heterotypic interaction has induced HSPC polarization with the formation of magnupodium and lysosomes clustering toward the site of contact, the subsequent division is marked by perpendicular spindle positioning, maintenance of the asymmetric distribution of lysosomes and final asymmetric inheritance of lysosomes in the siblings.

### Osteoblast interaction-driven HSPC asymmetric division boosts siblings heterogeneity

Lysosomes inheritance has been shown in human HSPCs to be predictive of cell surface markers expression in siblings. In particular, CD34 expression was found to be decreased in the daughter cell inheriting more mother cell lysosomes ^36^. To further characterize the daughter cells generated, we investigated the expression of cell surface markers of interest. Fluorescently labelled CD34 and CD33 antibodies were added in the microwells after the HSPC division (Figure 5A, Figure S3A and B). The distribution of CD34 and CD33 in daughter cells was quantified. For both markers asymmetric or symmetric distributions could be observed (Figure 5B and C). While CD33 sister ratios were similar in both conditions (Figure 5D), CD34 sister ratio was significantly higher upon culture with osteoblast than on fibronectin (Figure 5E).

**Figure 5.**
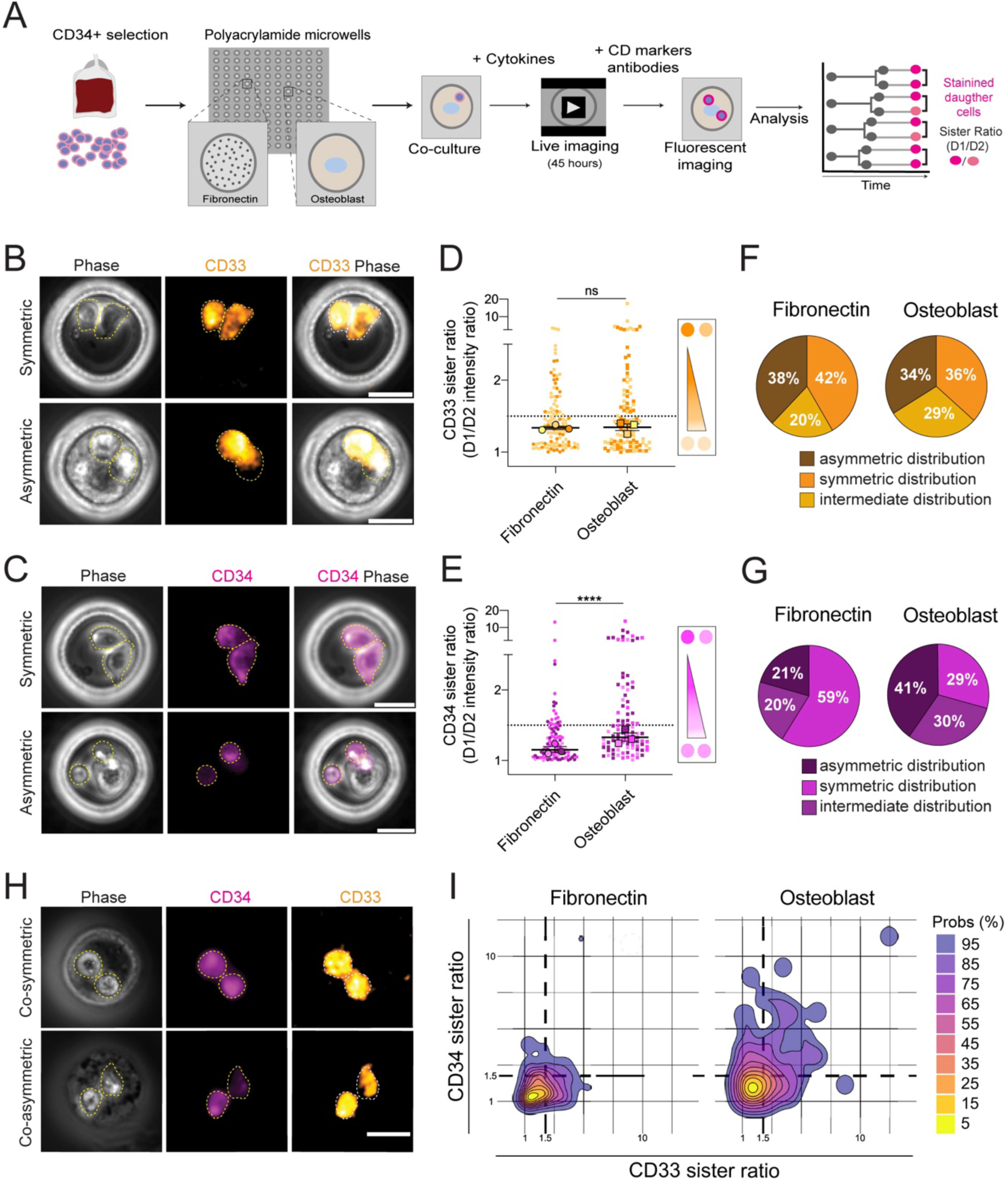
Osteoblast interaction fosters sibling heterogeneity. **(A)** Schematic representation of the experimental design for CD markers labelling and analysis of HSPCs siblings. **(B)** Representative transmitted light and fluorescence time-lapse images of HSPC labelled with CD33-PE, exhibiting symmetric (upper row) and asymmetric (lower row) distributions of the CD markers in the siblings **(C)** Representative transmitted light and fluorescence time-lapse images of HSPC labelled with CD34-APC, exhibiting symmetric (upper row) and asymmetric (lower row) distributions of the CD markers in the siblings. Daughter cells are contoured with yellow dashed lines. Scale bar: 20 µm. **(D)** SuperPlot analysis of CD33 marker signal ratios in daughter cells (sister ratio) in contact with fibronectin, and osteoblast (three biological replicates; fibronectin n_tot_=109, osteoblasts n_tot_=108 cells) **(E)** SuperPlot analysis of CD34 marker signal ratios in daughter cells in contact with fibronectin, and osteoblast (three biological replicates; fibronectin n_tot_=107, osteoblasts n_tot_=111cells). In D and E, each color represents a biological replicate. For each replicate, the median value appears as a large disk of the corresponding color. Mean of medians ± SEM shown as black bars. Kruskal–Wallis ANOVA test. ns: non-significant; ****: P≤0.0001. **(F)** Pie chart representation of CD33 distribution in HSPC siblings upon culture on fibronectin (left chart) and osteoblast (right chart). **(G)** Pie chart representation of CD34 distribution in HSPC siblings upon culture on fibronectin (left chart) and osteoblast (right chart). In F and G, CD markers distributions are classified into asymmetric (>1.5: dark purple for CD34 and dark brown for CD33), intermediate (≥ 1.25 and ≤1.5: magenta for CD34 and brown for CD33) and symmetric distributions (≤ 1.25: light magenta for CD34 and orange for CD33). **(H)** Representative transmitted light and fluorescence time-lapse images of HSPCs co-labelled with CD34-APC (magenta) and CD33-PE (orange) exhibiting symmetric (upper row) and asymmetric (lower row) co-distributions. Daughter cells are contoured with yellow dashed lines. Scale bar: 20 µm. **(I)** Density map representation of the CD34 versus CD33 ratios in siblings produced upon HSPC division in fibronectin or osteoblast conditions (three biological replicates, fibronectin n_tot_=100, osteoblast n_tot_=90 cells). The color code scale used for the probability representation is depicted on the right.

To further classify the daughter cell doublets, we set a sister ratio higher than 1.5 as indicative of an asymmetric distribution and a sister ratio lower than 1.25 of a symmetric distribution. In the case of CD33, the distribution of the populations was not significantly different in the two experimental conditions, with a third of the siblings exhibiting an asymmetric distribution (Figure 5F). In the case of CD34, asymmetric distribution was twice higher (41%) upon culture with osteoblast compared to fibronectin (Figure 5G). The distribution of both markers in siblings was analyzed (Figure 5H). The density maps generated in the two experimental conditions shed light on the differences in sibling heterogeneity landscapes. The whole population of siblings obtained on fibronectin covers exclusively 75% of the population obtained on osteoblasts. In the latter case, 25 % of siblings emerge as an original population highly divergent (Figure 5I).

These results indicate that heterotypic interaction with osteoblasts increases the level of CD34 asymmetric distribution in the daughter cells and in turn boosts the heterogeneity of the progeny.

## Discussion

We here demonstrate that heterotypic interaction is a genuine external cue which drives human HSPC asymmetric division using this model. This asymmetric division relies on a central role of the centrosome, a conserved feature of asymmetric divisions in solid tissues ^6^ and of lymphocytes engaged in immune synapse ^37 38^.

In HSPC, centrosome positioning in close vicinity of the contact site is associated with Golgi and lysosomes clustering at the cell periphery, as in lymphocytes engaged in immune synapse ^39^. Nonetheless, while lymphocytes tend to spread on their target cell, HCSPC interact with the osteoblast through the elongated magnupodium ^32^. This striking structural difference can be correlated to the half-lives of interaction in both systems: minutes in most of the cases for the immune synapse ^40^, hours in the case of HSPC. One may speculate that magnupodium assembly is an efficient strategy developed by the small sized HSPCs to sustain a robust and long-term polarization.

Interestingly, a fraction of HSPCs cultured in absence of polarizing external cues (fibronectin or skin fibroblasts) underwent asymmetric divisions. HSPCs therefore possess intrinsic properties that make them prone to asymmetry ^13 15^. However, heterotypic interaction with osteoblast, most likely through the magnupodium-associated hyperpolarization, had both a quantitative and qualitative effect on the division by creating a bias toward asymmetry, and generating siblings more drastically divergent. In addition, interaction with osteoblast gave rise daughter cells positioned proximally and distally to the osteoblast. The proximal cell inherited most of the mother lysosomes, indicative of a reduced commitment to differentiation ^41 42 36^. Such spatial control of the division is a conserved feature of asymmetric divisions in other stem cell niches ^6^ and may account for *in vivo* observations of organization into clusters of blood cells differentiation ^43^.

Stem cell asymmetric division is considered as a binary process leading to the generation of daughter cells of distinct identities ^44 45^. The early steps of hematopoiesis are now considered to occur like a continuum ^46^ with the slow and progressive emergence of distinct cell populations ^47^. Our data support the idea that heterotypic interaction driving HSPC asymmetric division generated siblings with different differentiation potentialities rather that distinct identities. Such non-stereotypical asymmetric divisions are likely to be a key process maintaining the plasticity of early steps of hematopoiesis.

## Materials and methods

### Mold fabrication for microwells

The mold design and fabrication were performed as previously described ^32^: microwell shape, size, and arrangement were drawn using the software CleWin and etched in the chrome layer onto a quartz photomask (Toppan Photomask). A wafer with microstructures was made on silicon. Silicon wafers were coated with a first 5-μm layer of resin (SU8-3005; MicroChem; CTS) exposed to UV light at 23 mJ/cm2 (UV KUB2; Kloe) for 5 seconds, and with a second layer, 50 μm thick (SU8-3050; MicroChem; CTS) exposed to UV light through the quartz mask at 23 mJ/cm2 for 8 seconds to create microwells. After development (Developer SU8; MicroChem; CTS), only the exposed structures remained, it was baked at 150°C for 2 hours and coated with gas-phase trichloro(perfluorooctyl)silane (Sigma-Aldrich). A negative mold of the silicon wafer was created using PDMS and treated with silane in the same way as the wafer. A second, positive mold of PDMS was made from the first mold, which will be referred to as the PDMS stamp from now on.

### Microwells fabrication

Clean glass coverslips were coated with a solution containing 3-(trimethoxysilyl) propylmethacrylate (Sigma-Aldrich), acetone, and ethanol, in a ratio of 1:0.5:50 for 10 minutes, followed by baking at 120°C for 1 hour. The PDMS stamp underwent plasma treatment for 30 seconds and was immediately placed on the coverslip. A freshly prepared solution containing 20% 37.5/1 acrylamide/bisacrylamide (Euromedex), in MilliQ water, along with 1% ammonium persulfate, tetramethylethylenediamine (Sigma-Aldrich), and 1% photo initiator (2-hydroxy-2-methylpropiophenone; Sigma-Aldrich), was introduced between the PDMS stamp and the glass coverslip using capillary action. The sample was exposed to UV light at a dosage of 23 mJ/cm2 for 5 minutes (Figure S1B). After exposure, the PDMS stamp was removed in MilliQ water and washed with 70% ethanol under UV light for 1 hour for sterilization. The samples were washed in PBS overnight to remove any remaining traces of toxic compounds.

### Cells and culture in microwells

All human umbilical cord blood samples collected from normal full term deliveries were obtained after mothers’ written and informed consent, following the Helsinki’s Declaration and Health Authorities. Human umbilical cord blood samples were obtained from the Cord Blood Bank of the Saint-Louis Hospital by French national law (Bioethics Law 2011-814) and under declaration to the French Ministry of Research and Higher Studies.

Mononuclear cells were collected using Ficoll separation medium (Eurobio, Courtaboeuf, France). CD34+ cells were further selected using Miltenyi Magnetically Activated Cell Sorting (MACS) columns (Miltenyi Biotech, Paris, France) according to the manufacturer’s instructions. CD34+ cells were then either put in culture or frozen at 80°C in IMDM medium (Gibco) supplemented with 10% fetal bovine serum (FBS) and 10% DMSO (WAK Chemie Medical GmbH).

The cell lines used were hFOB (osteoblasts, CRL-11372; ATCC) and BJ (skin fibroblasts, CRL-2522; ATCC), both cultured in DMEM-F12 medium (Gibco) supplemented with 10% FBS and 1% antibiotics (AA; Gibco, ref.15240062).

Microwells bottom glasses were coated with 10 μg/ml of fibronectin diluted in PBS for 5 minutes at 37°C. Microwells were gently rinsed twice in PBS, and a droplet of 40µL of supplemented DMEM F-12 was placed on top of the microwells. 5.000 osteoblasts or skin fibroblasts in 5µL volume were directly added to the DMEM-F12 droplet in a zigzag manner. After a 30-minutes incubation period to allow the seeding of osteoblast and skin fibroblast, the dishes were filled with DMEM-F12. Following a two-hour interval to facilitate cell spreading, 5.000 HSPCs were directly seeded onto the microwells using the same method. After a 30-minute incubation period to enable HSPC seeding, IMDM supplemented with 10% FBS, 1% AA, 100ng/mL human SCF (Peprotech, ref.300-07), 20ng/mL human IL-3 (Peprotech ref.300-23) and 10ng/mL human G-CSF (Prepotech ref.200-02) was added to mark the starting point (t0) of the experiment (Figure S1C).

### Cell labelling for live imaging

Coverslips with the microwells were pasted to the bottom of a bottom-left 35mm plastic dish, before the sterilization step. The medium was constantly changed every 24 hours.

For live imaging labelling, HSPCs were incubated with 20µM NBD C6-Ceramide (6-((N-(7-Nitrobenz-2-Oxa-1,3-Diazol-4-yl)amino)hexanoyl)Sphingosine) (Invitrogen) for 30 minutes at 4°C for Golgi staining or Cytopainter (Abcam) for Golgi and Nucleus staining. HSPC were incubated with 2µM Orange Cell-Tracker (ThermoFisher-LifeTech, ref.C34551) for 30 min or/and 1:250 LysoBrite Deep Red (Interchim, ref. 22646) or LysoBrite NIR (Interchim, ref.22641) for 50 min, both at 37°C. HSPCs were rinsed and pelleted twice with HBSS (Gibco) 10mM HEPES (Fisher). Labelled-HSPCs were finally washed in supplemented IMDM with 10mM HEPES and, after centrifugation, seeded in the microwells and cultured in phenol-red-free IMDM (Fisher) 10mM HEPES with cytokines, FBS, and AA. Images were taken every 30 minutes for transillumination and every 2 hours for laser illumination.

### CD staining of daughter cells

HSPCs were imaged using transillumination for 46 hours. After 46 hours and following the peak f cell divisions, anti-CD33-PE (555450, BD Bioscience) and anti-CD34-APC (55824, BD Bioscience) antibodies were added in a 1:60 ratio in 0.05% fetal human serum and PBS with 2mM EDTA for 20 minutes in the microwells.

### Immunofluorescence

Cells were fixed in PBS with 4% PFA for 15 min at 37°C after 20h or 40h of culture. Cells were permeabilized in 0.1% Triton X-100 for 20 minutes. Coverslips were neutralized with a solution of NH_4_Cl for 10 min. The following primary antibodies and dilutions were used: rabbit anti-pericentrin (ab4448; Abcam), 1/500; human anti-Giantin (A-R-H#03 TA10 hFc; Institut Curie), 1/200. Primary antibodies were incubated for 1 hour. The following secondary antibodies and dilutions were used: Alexa Fluor 568- and 647-conjugated goat anti-human and anti-rabbit (Life Technologies), 1:1000. Alexa Fluor 488-conjugated phalloidin (Sigma-Aldrich) was used to label F-actin. Finally, cells were incubated with DAPI (Sigma-Aldrich) for 10 min to stain the nucleus. The coverslips were mounted with Mowiol (Sigma-Aldrich).

### Microscopy

For the measurement of polarization index and mitotic spindle orientation, images were taken using a Nikon Ti-eclipse microscope equipped with a spinning disk (Yokogawa-CSU-X1) with a 60×, 1.5× 1.4-NA oil objective on an electron-multiplying charge-coupled device camera (Photometrics-Evolve512). MethaMorph was used as acquisition software. Z-stacks of 0.5µm were taken and a binning of 2×2 was used.

For live-cell imaging with trans and laser illumination light, an Olympus IX83 microscope equipped with a PECON CellVivo incubation system controlling temperature (37°C) and CO2 concentration (5%) was used. Images were acquired with a 20× 0.30-NA air objective on an ORCA-Flash4.0 Lite (Hamamatsu) camera and using MicroManager 1.4.21 software. A binning of 2×2 was used during the acquisition.

For detailed live-cell imaging of lysosome polarization and asymmetric inheritance with trans and laser illumination lights (Figure 3F, Figure 4, Figure S2B and C), a Nikon Ti2 Eclipse equipped with a Prime BSI Express CMOS camera (Photometrics) and using a Nikon CFI Plan Fluor 60x/0.75 NA oil objective with 1.5 objective amplification. PECON CellVivo incubation system was used to control temperature (37°C) and CO2 concentration (5%). Images were acquired with an ORCA-Flash4.0 Lite (Hamamatsu) camera and using MicroManager 1.4.21 software. To improve the signal/noise ratio of LysoBrite and reduce laser intensity and exposition time, HSPCs were pre-stimulated for 20h with the cytokines cocktail, then labelled with LysoBrite, and added to the microwells with osteoblast, as previously described. After 15 hours (35 hours of cytokines stimulation), microwells were positioned at the microscope and imaged every 15 minutes with ten distant 4µm Z-stack and 2×2 binning.

### Immunofluorescence quantification and analysis

#### Polarization Index, Golgi polarization, prophase centrosome positioning, and magnupodium

In the 3D images of the HSPCs, the positions of the centrosome A (Xa, Ya, Za), the point of contact B (Xb, Yb, Zb), and the most distal point on the HSPC membrane from the point of contact C (Xc, Yc, Zc), excluding thin membrane protrusions, were determined. The polarization index was calculated as d/D using ImageJ software. Distance D was defined as the length between points B and C, whereas distance d as the length between B and the projection of A in the D vector, using the following equation:

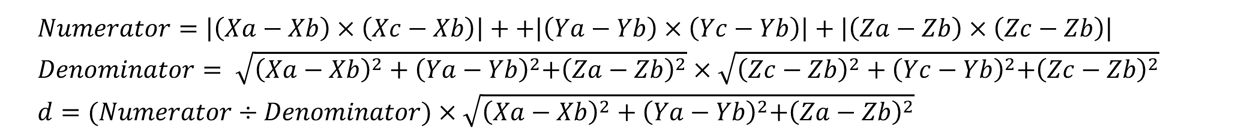

The same method was used for Golgi polarization analysis (taking the closest point of the Golgi to the cell membrane as “B”) and for both prophase centrosomes polarization analysis (polarization index ≤ 0.3 was taken as a proximal centrosome and > 0.3 as a distal centrosome). For magnupodium, the structure must be an elongated protrusion (longer than 2µm) with a close centrosome to be considered. For the data presented as SuperPlots ^48^, each color represents a biological replicate, and the medians for each replicate, the mean of the medians, and the standard error of the mean are presented. SuperPlots were only used in the cases where each biological replicate had more than 20 cells. The normality of data distribution was tested by Shapiro–Wilk test for further statistical test selection. The significance of the difference between populations was tested using nonparametric (Kruskal– Wallis) ANOVA and Mann–Whitney t-test for no-Normal distribution samples, or parametric unpaired t-test and One-way ANOVA for Normal distribution samples. All statistical analyses were performed with Prism software (GraphPad).

#### Mitotic Spindle Orientation

Mitotic cell images were resliced to visualize the centrosomes and the point of interaction with the bottom of the microwell or the osteoblast/fibroblast. For the mitotic spindle angle, the angle generated between both centrosomes with respect to the image X axis was subtracted to the angle generated for the point of interaction with respect to the image X axis. Graphs were generated with R ^49^.

### Live imaging quantification and analysis

#### Live Golgi and lysosome polarization

Line-scan from the contact site to the more distal point was used to assess the lysosome localization with respect to the Golgi in the Max projected images. Graphs were done with Prism software (GraphPad).

#### Lysosome inheritance ratio

cells were tracked manually until division. To automatize the daughter cell area selection, CellTracker images were processed with Gaussian Blur (sigma=2), Subtract Background (rolling=50), and then thresholding (Otsu dark) to create a mask for the daughter cell area. Background signal was selected manually inside the microwell for each image. Lysosome inheritance ratios were calculated in untreated images using the first time point following cell division, by dividing the brightest daughter cell by its sister, using the sum of LysoBrite pixel fluorescence values (CTCF) normalized to the CellTracker CTCF for each cell. Ratios were therefore ≥ 1. Ratios <1.5 and ≥ 1.5 were respectively selected as indicative of symmetric and asymmetric lysosome inheritance. In the cases where both daughter cells were very close in the first time point after division, the following time point was analyzed. Manual selection of the daughter area was done when the automatized selection failed. Data were analyzed with Fiji and presented as SuperPlots in the same way previously referred. All statistical analyses were performed with Prism software (GraphPad).

#### Lysosome polarization, mitotic spindle orientation, and inheritance ratio

Lysosome polarization of mother cells were analyzed as previously described in the time-lapse before chromosome condensation, visualized by phase images. Metaphase/anaphase was analyzed in the time-lapse that allow the visualization of the chromosome orientation to determine mitotic spindle orientation with respect to the interaction point. In the cases where the chromosome orientation was not clear, the visualization of a cell-side attachment from metaphase to telophase was taken as a perpendicular division. Lysosome inheritance was measured after the division in the first-time frame that better visualize the whole volume of the two daughter cells.

#### CD marker ratios

Gaussian fit curves were created based on the CTCF values of non-staining and staining HSPCs for the two CD marker signals. 95% confidence intervals of the non-staining intestines were selected to determine the thresholds for positive CD34 and CD33 signals. Single mother cells were tracked manually until their division by transillumination to annotate the division time. To automatize the daughter cell area selection, CD34 fluorescence images were processed with Gaussian Blur (sigma=2), Subtract Background (rolling=50) and then thresholding (Otsu dark) to create a mask for the daughter cell area. The background region was selected manually inside the microwell for each microwell. When the CD marker CTCF of both daughter cells was below the threshold, the cells were discarded. When one daughter cell was below the threshold, its CTCF was automatically converted to the value threshold and both sister cells were compared. CD marker production ratios were calculated after antibody incubation, by dividing the brightest daughter cell by its sister. Ratios were therefore always ≥ 1. Ratios ≤ 1.2 and ≥ 1.5 were respectively selected as indicative of symmetric and asymmetric distributions. Ratios between 1.2 and 1.5 were indicative of intermediate distribution. In the case where both daughter cells were overlapping, the next time point was analyzed. Manual selection of the daughter area was done when the automatized selection failed. Data were analyzed with Fiji and presented as SuperPlots, or with R ^49^. In Figure 5, the corresponding density map was generated using the “ggdensity” package ^50^. Statistical analyses were performed with Prism software (GraphPad).

## Supplemental Figures

**Supplemental Figure 1.**
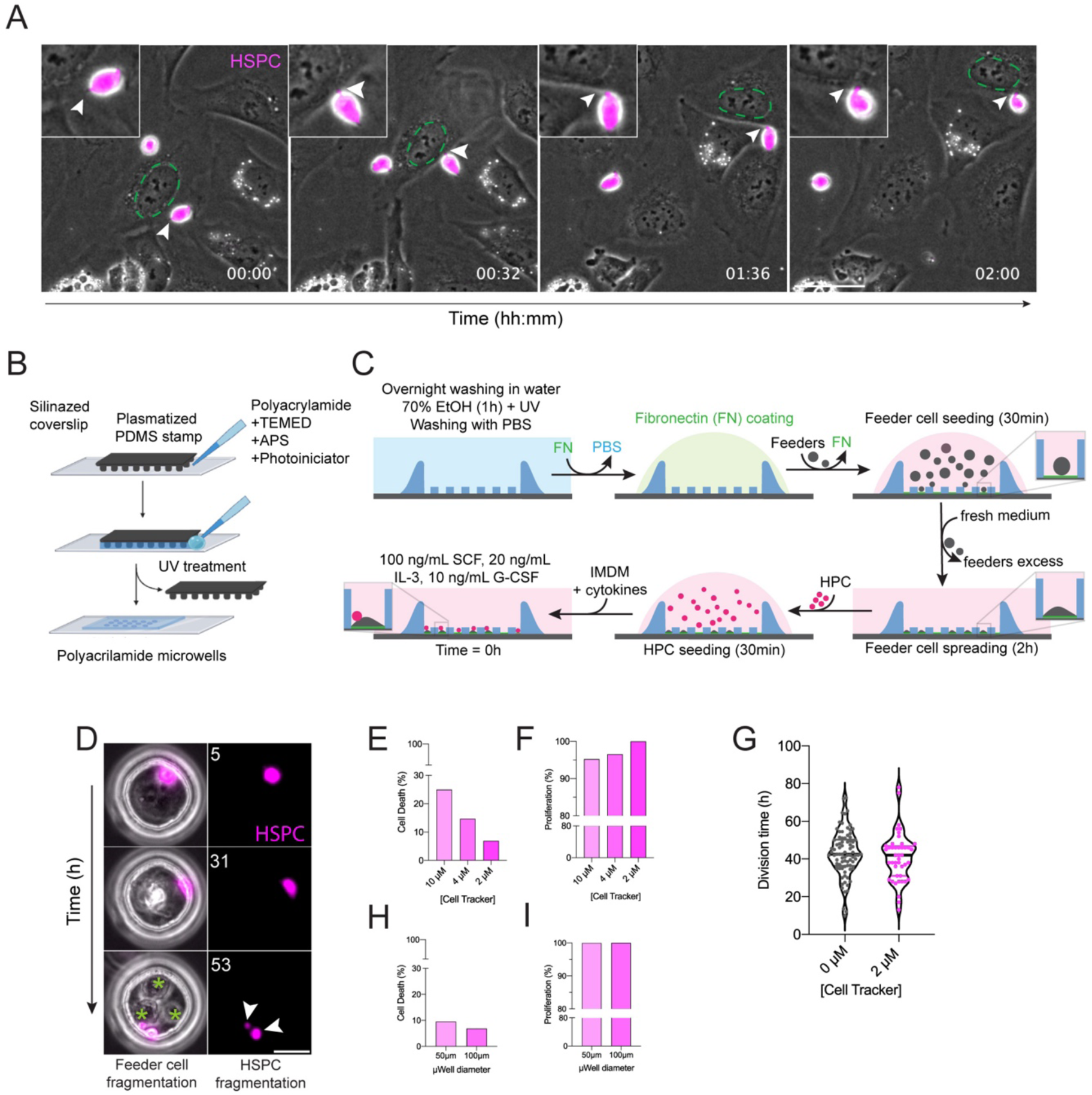
Microwells as an optimal experimental system to assess HSPCs cell divisions upon heterotypic interaction. **(A)** Time-lapse monitoring with transmitted light and fluorescence of live HSPCs labelled with CellTracker (magenta) in contact with osteoblasts in a classical 2D co-culture system. Green circle marks the nucleus of interacting osteoblast over time. Arrows highlight the site of contact of HSPC with the migrating osteoblast. Zoom images centered on the interacting HSPC are presented as inserts in the up-left corners. Time is indicated in hours: minutes. Scale bar: 20µm. **(B)** Scheme of polyacrylamide microwell fabrication using a PDMS mold. **(C)** Scheme of the cell seeding protocol based on first fibronectin coating, then feeder cell seeding, and finally, the addition of the HSPCs. **(D)** Time-lapse monitoring with transmitted light and fluorescence of live HSPCs labelled with CellTracker (magenta) in contact with osteoblast. Osteoblast fragmentation is observed (green asterisks) and HSPC cell death (arrows). Time is indicated in hours. Scale bar: 20µm **(E)** Quantification of HSPC cell death according to the CellTracker concentration and upon co-culture with osteoblast in microwells of 100µm diameter (one biological replicate,10µM n_tot_=28, 4 µM n_tot_=34, 2µM n_tot_=29 cells). **(F)** Quantification of HSPC proliferation according to the CellTracker concentration and upon co-culture with osteoblast in microwells of 100µm diameter (one biological replicate, 10µM n_tot_=21, 4 µM, n_tot_=29, 2µM n_tot_=27 cells). **(G)** HSPC timing of division in absence or presence (n_tot_=82 and 44 cells) of CellTracker and fluorescence light. HSPC were co-cultured with osteoblasts in microwells of 50µm diameter (two biological replicates). **(H)** Quantification of HSPC death according to the microwell diameter. HSPC were co-cultured with osteoblast and stained with 2µM of CellTracker (one biological replicate, 50µm diameter n_tot_ = 21, 100µm diameter n_tot_=29 cells). **(I)** Quantification of HSPC proliferation according to the microwell diameter. HSPC were co-cultured with osteoblast and stained with 2µM of CellTracker (one biological replicate, 50µm diameter n_tot_=19, 100µm diameter n_tot_=27 cells).

**Supplemental Figure 2.**
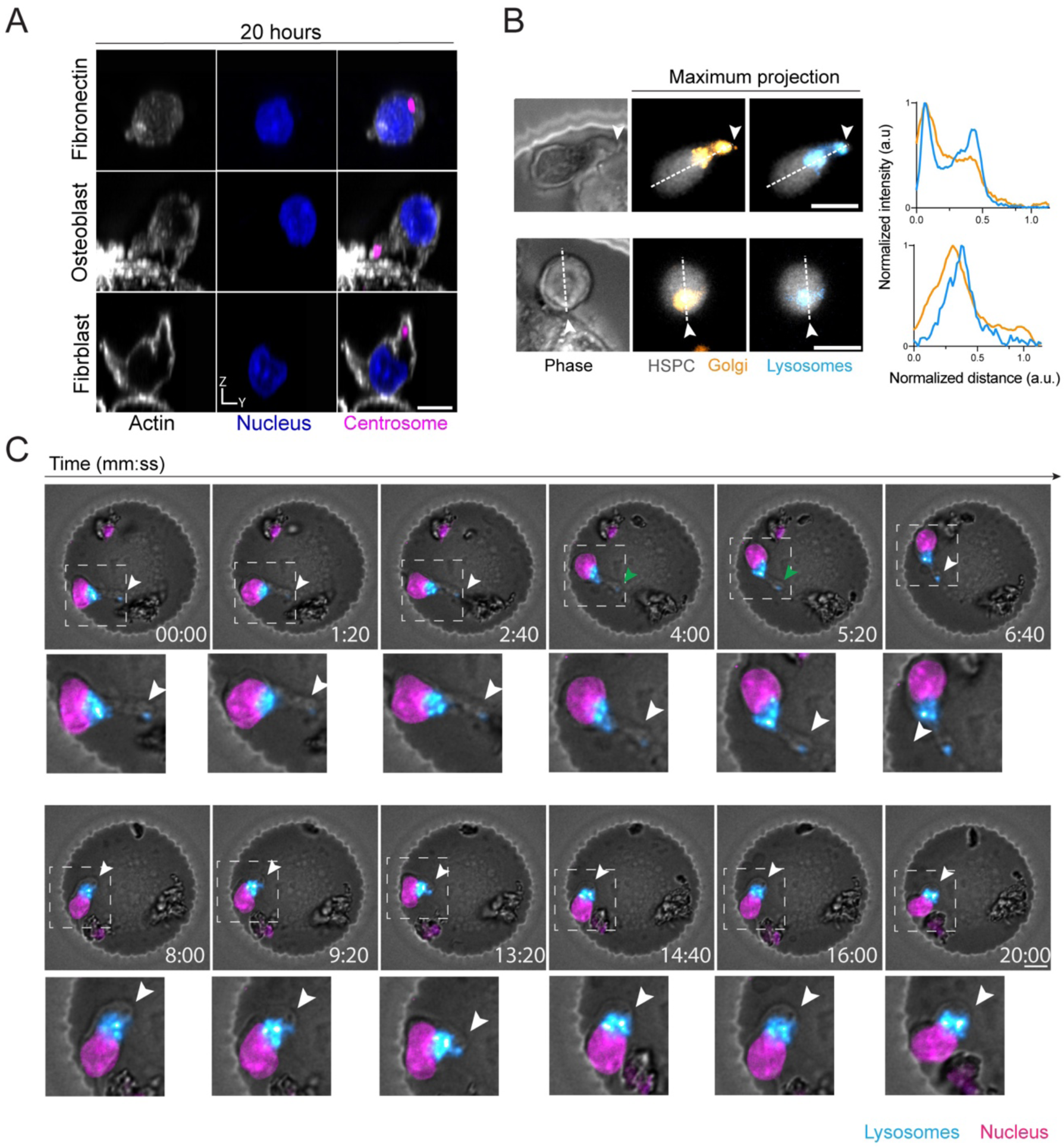
HSPC polarization after 20h of culture. **(A)** Representative images of HSPC in contact with fibronectin, osteoblasts, and skin fibroblasts after 20 h of culture. Actin appears in white, the nucleus in blue, and the centrosome in magenta. Scale bar: 5µm. **(B)** Representative transmitted light and fluorescence time-lapse images of live HSPCs in contact with osteoblast. HSPC is labelled with CellTracker (gray), Golgi tracker (orange), and LysoBrite (blue) for lysosome live staining. Scale bar: 5 µm. Associated line-scan analysis for Golgi (orange) and Lysosome (blue) signal intensity is shown on the right of the analyzed images. **(C)** Time-lapse monitoring with transmitted light and fluorescence of live HSPCs labelled with nucleus tracker (magenta) and LysoBrite (blue). HSPC interaction is stable in time and marked by lysosomes enrichment toward the site of interaction. Time is indicated in minutes and seconds. Scale bar: 20µm. Zoom in on the HSPC interaction with the osteoblast is shown below. Arrows highlight the site of contact of the HSPC with the osteoblast.

**Supplemental Figure 3.**
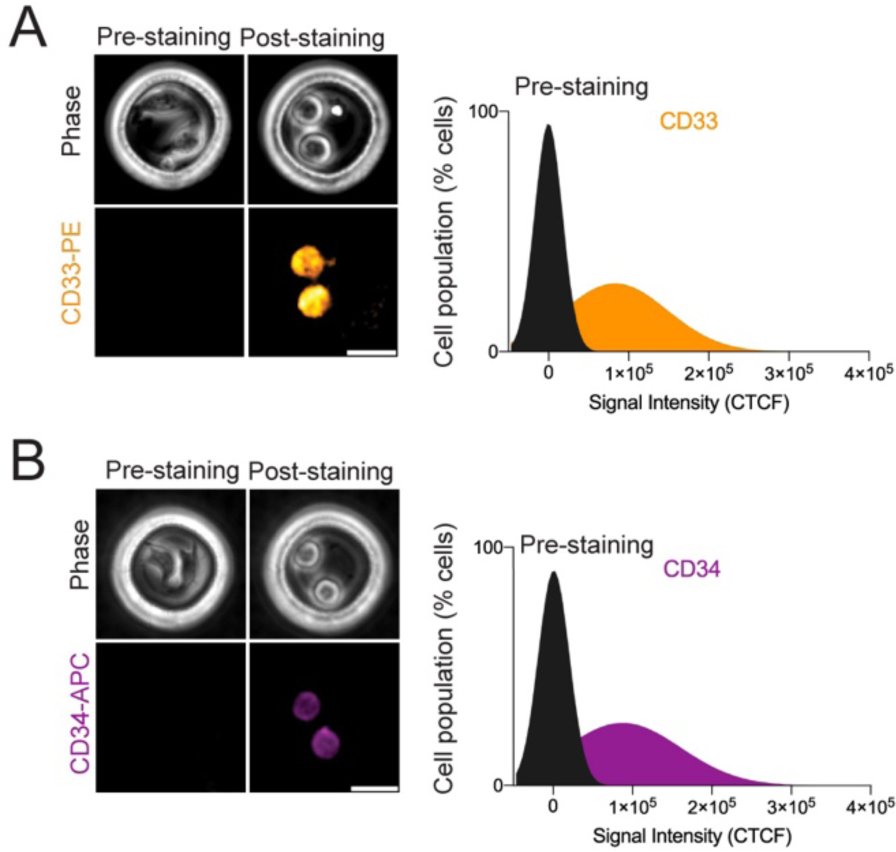
CD marker signal and effect of the daughter cell cycle age in CD markers production. **(A)** Left panel: representative transmitted light and fluorescence time-lapse images of HSPC before and after labelling following cell division with CD33-PE. Scale bar= 20µm. Right panel: analysis of CD33 signal intensity (CTCF) before labelling and in labelled daughter cells (two biological replicates, unstained n_tot_=124, stained cells n_tot_=238 cells). **(B)** Right panel: representative transmitted light and fluorescence time-lapse images of HSPC labelled with CD34-APC, Scale bar= 20µm. Right panel: analysis of CD34 signal intensity (CTCF) before labelling and in labelled daughter cells (two biological replicates, unstained n_tot_=124, stained cells n_tot_=238 cells).

## Acknowledgements

This work was supported by the European Research Council (Consolidator Grant 771599 (ICEBERG) to MT and Advanced Grant 741773 (AAA) to LB), by the Bettencourt-Schueller foundation, the Emergence program of the Ville de Paris and the Schlumberger foundation for education and research. AC received PhD fellowship from the Fondation pour la Recherche Médicale (Grant FDT20170437071). We thank the Technological Core Facility Research Institute, UMS “Saint-Louis”, US53/UAR2030. The facility is supported by grants from Université Paris Cité, ITMO-Cancer of the French National Science Alliance (Aviesan), the Conseil Regional d’Ile-de-France (Canceropôle), the National Cancer Research Institute (InCa), the Ministère de la Recherche, the Association Saint-Louis and the Association Jean Bernard. We thank Sandrine Moutel and the platform TabIP of the Curie Institute (Paris) for sharing the human anti-Giantin antibody.

## Author contributions

A.C performed all the experiments, with the help of S.B and L.F for HSPC collection and processing. A.C performed all the data analyses, with the help of M.G for data filtering and representation using R. B.V was involved in the development of technological implementations. S.B, M.T, J.L and L.B supervised the work and obtained funding for the project. A.C and S.B wrote the manuscript with the help of M.T, the manuscript was further critically reviewed by all authors.

## Inclusion and diversity

We support inclusive, diverse, and equitable conduct od research

